# Disconnection Between Microvascular Damage and Neurodegeneration in Early Diabetic Retinopathy

**DOI:** 10.1101/2024.01.31.577770

**Authors:** Qian Yang, Marina Yasvoina, Abraham Olvera-Barrios, Joel Mendes, Meidong Zhu, Cathy Egan, Adnan Tufail, Marcus Fruttiger

## Abstract

**Aim:** Diabetic retinopathy (DR) is a common complication of diabetes mellitus and can result in vision loss. Early clinically diagnosed signs of DR are linked to vascular damage, impacting on the neural retina typically at later stages. However, vascular changes and potential effects on neural cells before clinical diagnosis of DR are less understood.

**Methods:** To learn more about the earliest stages of DR we studied postmortem retina from diabetic donors who did not have clinical DR. Histological phenotyping and quantitative analysis were carried out on retina from 14 donors (3 controls, 10 diabetics, and 1 DR case) to examine capillary loss in the deeper vascular plexus (DVP) and the superficial vascular plexus (SVP) of the retinal vasculature and to study effects on the neural retina.

**Results:** The advanced DR case exhibited profound vascular and neural damage as expected, whereas none of the ten randomly selected diabetic donors had any DR signs that would have been diagnosed clinically. Within that group, two showed very minor capillary dropout in the SVP, whilst in the remaining diabetic cases the SVP was indistinguishable from the controls. In contrast, over half of the diabetic retinas showed capillary dropout in the DVP and increased capillary diameter. Furthermore, a pan retinal loss of inner nuclear layer cells was observed in those diabetics with capillary dropout compared to the controls (p<0.05), but there was no local spatial correlation between neural cell loss and capillary dropout.

**Conclusions:** Our study has established a novel histological biomarker for diabetes related tissue damage at the earliest stages of DR in human postmortem retina, which appears to be common in people with diabetes before DR can be clinically diagnosed. Furthermore, the spatial mismatch between local capillary dropout and the diffuse neural loss in the INL suggests that at this very early stage of DR, microvascular loss may not causally be directly connected to neurodegeneration and that diabetes may affect the two readouts independently.

**RESEARCH IN CONTEXT:** What is already known about this subject?

- Early DR is currently clinically defined by visible microvascular changes.
- Tests of retinal function in people with diabetes imply neuroretinal impairments in early DR.
- Clinical imaging in people with diabetes suggests reduced perfusion in the deep retinal vasculature plexus.

What is the key question? (one bullet point only; formatted as a question)

- How does retinal microvascular damage relate to neuronal damage?

What are the new findings?

- We have developed a sensitive histological biomarker for microvascular damage in postmortem retinal tissue.
- Early vascular changes in the deeper plexus occur in diabetes even in the absence of manifest DR.
- Neurodegenerative changes are distributed throughout the retina in diabetes and do not spatially correlate with microvascular nonperfusion.

How might this impact on clinical practice in the foreseeable future?

- Deeper plexus perfusion can be a useful early biomarker to assess DR.

## INTRODUCTION

Diabetic retinopathy (DR) is a prevalent complication of diabetes and affects approximately one-third of people with diabetes globally [1]. One of the earliest recognised pathologies in DR is localised loss of small capillaries resulting from the death of endothelial cells and pericytes [2, 3]. This process gives rise to acellular capillaries, also known as string or ghost vessels, which are composed of the remaining vascular basement membrane [4]. These acellular vessels have been first described in human DR eyes over fifty years ago [5] and have also been extensively reported in various diabetes animal models [6–8]. They are not perfused [9, 10] and can therefore serve as a histological biomarker of non-perfusion postmortem.

In clinical practice, retinal vascular perfusion has been traditionally evaluated using fluorescent angiography (FA). However, the advent of optical coherence tomography angiography (OCTA) presents a less invasive alternative. OCTA also has the added advantage of detecting perfusion across all layers of the retinal vasculature, unlike FA, which is mainly limited to the superficial plexus and does not image the deeper capillary plexus well [11]. While several OCTA studies have identified reduced retinal vasculature perfusion in patients with diabetes, it remains a matter of debate whether the earliest abnormalities manifest in the deeper capillary plexus (DVP) or the superficial capillary plexus (SVP) (reviewed in **Supplementary material 1**).

Non-perfusion of the retinal vasculature typically leads to hypoxia and upregulation of vascular endothelial growth factor (VEGF) but can also cause neural cell death and retinal degeneration. Several studies have shown neural cell death and retinal tissue atrophy in patients with DR or retinal vascular occlusions [12–15]. However, the widely accepted view that neurodegenerative changes in DR are a secondary consequence of the primary vascular damage has been questioned [16–18]. There are several examples illustrating that the reverse (vascular dropout being secondary to neurodegeneration) can also occur. For instance, vessel density loss in the DVP has been observed in cases of retinal pigmentosa, which is well established to be primarily a neurodegenerative disease [19]. Furthermore, in patients with diabetes subtle functional changes, such as those observed in electroretinograms (ERGs), have been detected before the onset of noticeable damage to the retinal vasculature [20]. Regional variations in multifocal ERGs have also been found to predict future vascular lesions [21], suggesting that the earliest neuronal defects might occur independently of perfusion defects. Moreover, thinning of the nerve fibre layer has been demonstrated by OCT and histologically in people with diabetes and no to minimal DR [22, 23].

To investigate links between neural loss and capillary drop-out during the earliest stages of retinopathy in human tissue, we collected post-mortem eyes from anonymous donors with diabetes without clinical DR and quantified the number of acellular capillaries in their retinal vasculature. This allowed us to identify and study postmortem retinal tissue from donors with diabetes and with histological signs of early-stage DR, based on subtle microvascular defects, in the absence of a clinical diagnosis of diabetic retinopathy in the donors.

## RESEARCH DESIGN AND METHODS

### Donor Tissue Information

The study has ethical approval (UK National Research Ethics, IRAS project ID: 279162) and follows the tenets of the declaration of Helsinki. All donors documented their willingness to participate in the eye tissue donation programme of the study. A total of 14 human post-mortem eyes from 14 donors (10 donors with diabetes) were obtained from Moorfields Eye Bank (London, UK) and Lions NSW Eye Bank (Sydney, Australia), as shown in Error! Reference source not found.. Quality control on cross sections was performed for every eye analysed, and only those with normal appearing morphology were included for further investigations.

### Tissue Processing

The central region of the eye was dissected to include optic disk, fovea, nasal and temporal retinal periphery. Dissected samples were stained with rhodamine labelled Ulex Europaeus Agglutinin I (UEA, RL-1062, Vector Laboratories; 1:500) diluted in wholemount blocking buffer (1% FBS, 3% Triton X-100, 0.5% Tween 20 and 0.2% sodium azide in 2X PBS) overnight 4°C. After obtaining satisfactory wholemount images (see below), samples were washed in PBS overnight, dehydrated and paraffin embedded (LEICA TP1020 tissue processor, Leica, UK) and 6 µm sections were cut naso-temporally.

### Immunohistochemistry on Paraffin Sections

Staining was performed as previously described [24]. Heat induced antigen retrieval was performed using solution constituting of 90% glycerol and 10% 10Mm sodium citrate pH 6. Slides were heated in the solution to 120°C and kept for 20 min, then washed in ddH_2_O and incubated in blocking buffer (1% BSA, 0.5% Triton and 0.2% sodium azide in PBS) for 1 hour. Primary antibodies were diluted in blocking buffer and incubated at 4°C overnight, **Table 2** shows the details of primary antibodies in this study. After washes in PBS, sectioned were incubated in secondary antibody (1:200) for 1 hour at room temperature. Then slides were washed with PBS and counterstained with Hoechst (H33258, Sigma-Aldrich, St Louis, MO; diluted 1:10,000 in PBS) and mounted.

**Table 11.** Donor tissue and fixation information.

| Group | WM phenotype | Cross-section phenotype | Donor | Age | Sex | DM type | Cause of death | Fixation delay (h) |
| --- | --- | --- | --- | --- | --- | --- | --- | --- |
| Controls | No dropout | Controls | 1 | 69 | M | None | mesothelioma | 10 |
|  |  |  | 2 | 73 | F | None | end stage COPD | 19 |
|  |  |  | 3 | 67 | F | None | breast cancer | 22.5 |
| Diabetes with no diagnosed DR | No dropout | No dropout | 4 | 76 | M | II | metastatic gastric cancer | 12 |
|  |  |  | 5 | 87 | M | II | lung cancer | 28 |
|  |  |  | 6 | 62 | M | II | malignancy | 24.5 |
|  |  |  | 7 | 72 | M | N/A | lung adenocarcinoma | 20 |
|  | No dropout | Dropout | 8 | 55 | M | I | N/A (found unresponsive) | 18 |
|  | No dropout |  | 9 | 87 | M | II | myocardial infarction | 9 |
|  | Sporadic minor dropout |  | 10 | 74 | F | II | cardiac arrest | 8.75 |
|  | No dropout |  | 11 | 79 | F | II | metastatic ovarian cancer | 7.5 |
|  | No dropout |  | 12 | 66 | M | II | respiratory failure | 26.5 |
|  | Sporadic minor dropout |  | 13 | 61 | F | II | metastatic adenocarcinoma | 17 |
| DR |  | DR | 14 | 64 | F | II | myocardial infarction | 57 |

Abbreviation: I: Type I diabetes mellitus; II, Type II diabetes mellitus; WM, wholemount; COPD, chronic obstructive pulmonary disease; N/A, not applicable

**Table 2.** List of primary antibodies used in the study.

| Antibody | Company | Catalogue number | Dilution |
| --- | --- | --- | --- |
| Aquaporin-4 | Novus | NBP187679 | 1:300 |
| Collagen IV | Chemicon | AB769 | 1:500 |
| Collagen IV | BioRad | 2150-0140 | 1:500 |
| CRALBP | Thermo Fisher | MA1-813 | 1:300 |
| GFAP | Sigma | C9205 | 1:500 |
| Glutamine synthetase | Millipore | MAB302 | 1:300 |
| Parvalbumin | Swant | 235 | 1:500 |
| PKCα | Santa Cruz | sc-17769 | 1:500 |

### Microscopy

Retinal whole mounts stained with rhodamine-UEA were imaged using an Olympus SZX16 stereoscope (Tokyo, Japan) to generate an overview of the vasculature to evaluate vascular abnormality. Zeiss Axioskop 2 Upright Fluorescence Microscope (Carl Zeiss Microscopy LLC, USA) or Invitrogen™ EVOS™ FL Auto 2 (Thermo Fisher, UK) was used to visualize capillaries on cross sections. Regions of interest were then imaged with Zeiss LSM 700 with 40X objective lens. For consistency, six z-stacks with interval of 1 μm were taken for each region of interest.

### Image Processing and Analysis

Adobe Photoshop CS6 (Adobe Systems, Inc.) was used to generate whole mount panorama images with the built-in “Load Files into Stacks” and “Auto-Blend Layers” functions followed by manually stitched regions together. For cross sections, the built-in “Panorama” function was used. Manual alignment and adjustment were performed when needed.

### Quantification of Vessel Dropout

The entire dissected tissue was sampled by immunostaining three consecutive sections from every hundredth section. Retinal capillaries were firstly counted and assigned to a vasculature complex based on collagen IV and Hoechst staining. Endothelial lining was then identified independently based on UEA staining. A basement membrane (collagen IV+) without internal endothelial lining (UEA-) was considered as vessel loss, or ghost vessel [24]. Normal and ghost vessels in the different vascular plexuses were counted manually. Capillary dropout was expressed as the percentage of ghost vessels over total vessels overall or within the same plexus. Plexuses segmentation criteria followed what has been proposed previously [25]. Regions where the retinal vasculature plexuses cannot be distinguished, for instance, around the optic nerve head, were excluded from analysis.

### Nearest Neighbour Distance of DVP

To reveal the spacing profile of capillaries in the DVP, the nearest neighbour distance (NND) was measured, which has been a widely used method to quantify cell and capillary spacing profiles [26, 27]. In this study, we measured the distance to its nearest neighbour of each capillary residing in the inner nuclear layer (INL) on cross sections. This was performed manually using Fiji (NIH, RRID: SCR_002285). For each donor, NND was measured on every hundredth sections, at least five sections were examined, resulting in at least 1000 measurements for each donor. To correct for possible tissue shrinkage during processing, the height of the retinal pigment epithelium (RPE) nuclei was measured and used as a reference. At least 15 measurements were performed per donor. NND values of each donor were normalized to its reference accordingly. Data were presented as fold change to the reference.

### INL Cell Loss within Zone of Influence

On cross sections, a circle was drawn from the basement membrane of each capillary with the radius of half of the mean NND. Considering the upper and lower boundaries of INL, nuclei were counted in a circle around each DVP capillary. LSM700 laser scanning confocal microscopes (Zeiss) with 40X objective lens was used to visualize immunolabelled inner nuclear layer neurons. Five z-stacks of 1-µm steps, were scanned in each field analysed. The number of nuclei was counted manually. Random distribution simulation of the same sample size, mean and standard deviation was generated using Microsoft Excel.

### Data Analysis

Datasets were assessed for normal distribution using Shapiro-Wilk test with a significance level of 0.05. If datasets were normally distributed, two-tailed Student t-test was performed to compare the means of two groups. Otherwise, nonparametric Mann-Whitney test was used to compare the median and distribution of two datasets. One-way ANOVA test was used to compare the means of more than two groups, Dunnett’s post hoc analysis for comparing the means of control and other groups, and Tukey’s post hoc analysis for comparing the means of any two groups. P values in ANOVA test were adjusted. P-values below 0.05 were considered statistically significant. Because of the limited availability of postmortem DR tissue, we had only one cases diagnosed with DR. It served as a positive control, and the one sample t-test was used to compare values from the DR sample with mean values of other groups of interest. For linear regression, Pearson correlation coefficient was calculated to assess potential relationship between datasets. Linear relationship was assumed when the slope of the best fit model was significantly different from zero. All data are presented as mean ± standard deviation unless otherwise stated.

## RESULTS

### Loss of endothelial cells in retinal vasculature

To gain an overview of potential non-perfusion in the retinal vasculature of postmortem retina from donors with diabetes, we visualised endothelial cells in retinal wholemounts, using fluorescently labelled Ulex Europaeus Agglutinin I (UEA). **Figure 1A-C** is a representative sample of a whole mount image from a patient with type II diabetes. It shows small regions of endothelial cell loss in the peripheral retina (outside the standard 6X6mm screening area of OCTA, which would not usually be captured during standard clinical assessment).

**Figure 1.**
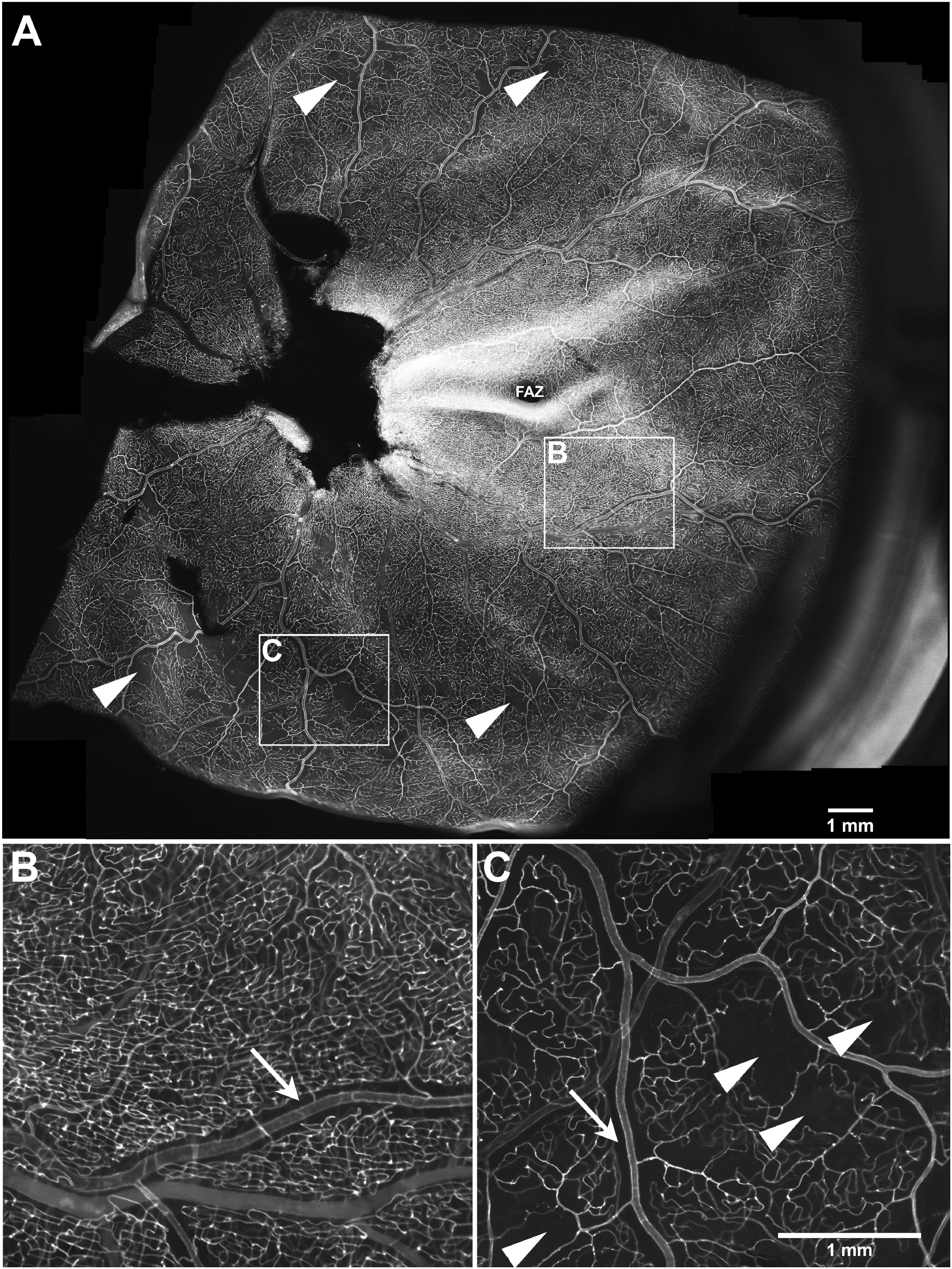
Endothelial cells in retinal whole mount. **A**, Example of a retinal whole mount stained with UEA revealing endothelial cells (optic disc and sclera have been removed). **B** and **C** are zoomed-in images of boxes shown in A. Capillary free regions can be seen in the vicinity of arteries (arrow in **B** and **C**), which is normal. In contrast, localised small capillary-free patches in the periphery (arrowheads in **A** and **C**) are indicative of retinal vasculature pathology. FAZ = foveal avascular zone. Scale bar 1mm.

A total of 8 out of 10 retinal whole mounts from donors with diabetes did not show any obvious loss of capillaries in this readout and appeared very similar to controls, **Table 1**). However, since it is difficult to directly measure the loss of individual capillaries (as they are no longer there), we developed a more sensitive method to measure damage in the retinal vasculature. A previous study from our group has shown that the basement membrane of non-perfused vessels is preserved for years despite the loss of the endothelial lining [24]. We therefore co-stained retinal sections with UAE and an antibody against collagen IV, visualising vascular basement membrane to identify acellular capillaries (**Figure 2 and Supplementary material 2)**, providing a quantifiable surrogate readout of loss of perfusion.

**Figure 2.**
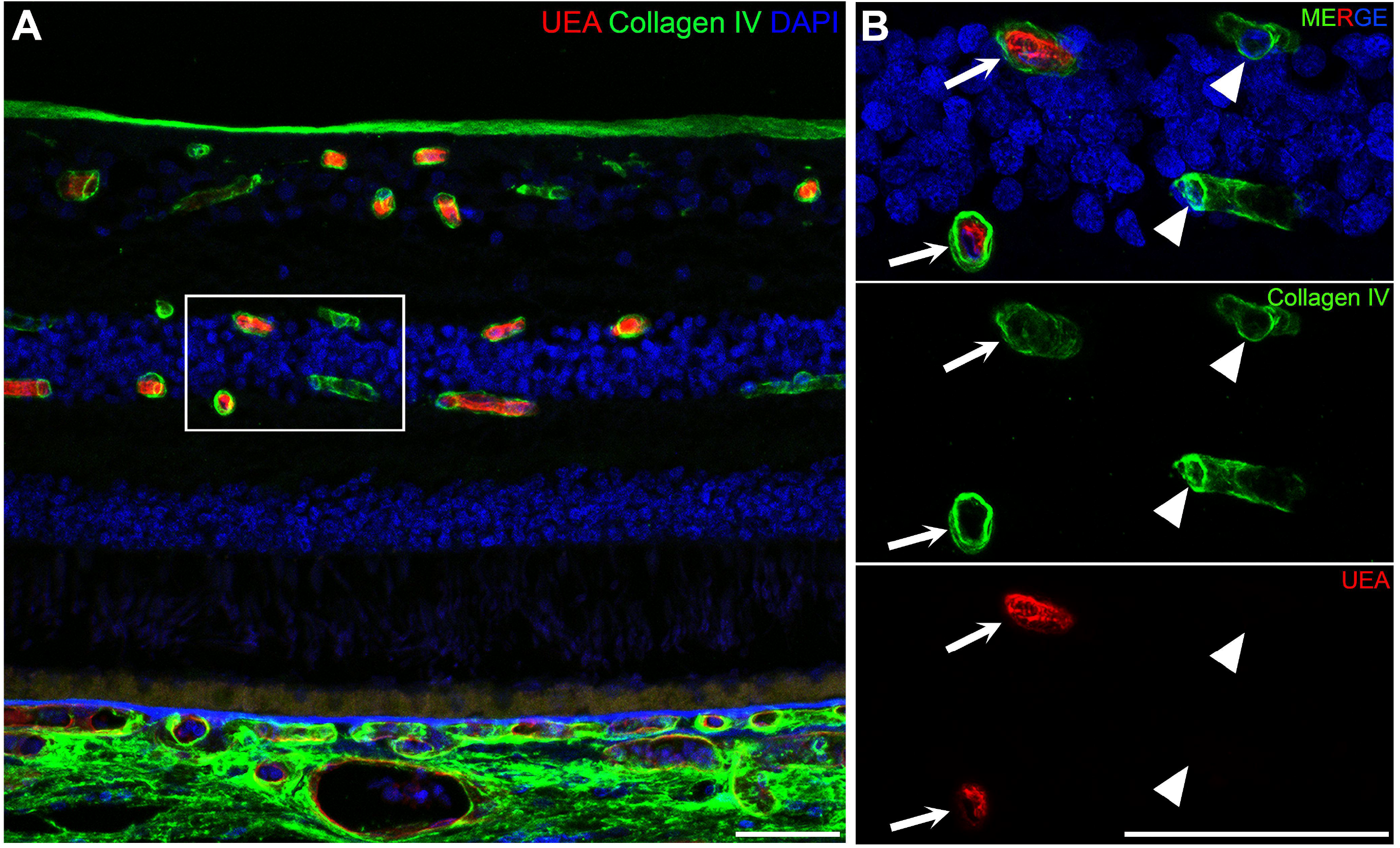
Immunostaining showing ghost vessel on a cross section from a diabetic eye. **A**, Immunohistochemistry showing UEA labelled endothelial cells and collagen IV-stained basement membrane from a region with acellular capillary of donor 10 (diabetic with dropout subgroup). Higher magnification of the boxed area in **A** is shown in **B**. Internal lining of endothelial cells was present in normal capillary (arrows), but not in ghost vessels (arrowheads). Scale bar 50 μm.

A total of 14 retinas (from 14 donors, **Table 1**) were chosen for quantitative analysis (3 controls, 10 patients with diabetes and no clinically diagnosed DR and 1 donor with DR). The retinas showed different amounts of acellular capillaries (**Fig. 3A**). Based on the frequency of acellular capillaries, two subgroups clearly emerged in the group with diabetes. The subgroup with diabetes and no dropout (DNDO) presented a level of capillary dropout (1.69 ± 0.37%) below the overall mean of the whole group with diabetes (3.82 ± 1.89%, **Figure 3A**, dashed line) and similar to controls. In contrast, the subgroup with diabetes and dropout (DDO), had a notably higher incidence of acellular capillaries (5.23 ± 0.57%). As assessed by one-way ANOVA test with *post hoc* analysis, the means of the control and the two subgroups with diabetes were significantly different from one another (p < 0.001). Whilst the DNDO subgroup appeared to have only slightly elevated degree of vessel loss compared to controls (p = 0.04), there was clearly more vessel loss in the DDO (p < 0.0001 when compared to controls or DNDO subgroup). Not surprisingly, in the retina with DR, the incidence of vessel loss was considerably increased (29.83%), indicating substantial vascular damage in advanced DR (p < 0.001 compared to any other groups).

**Figure 3.**
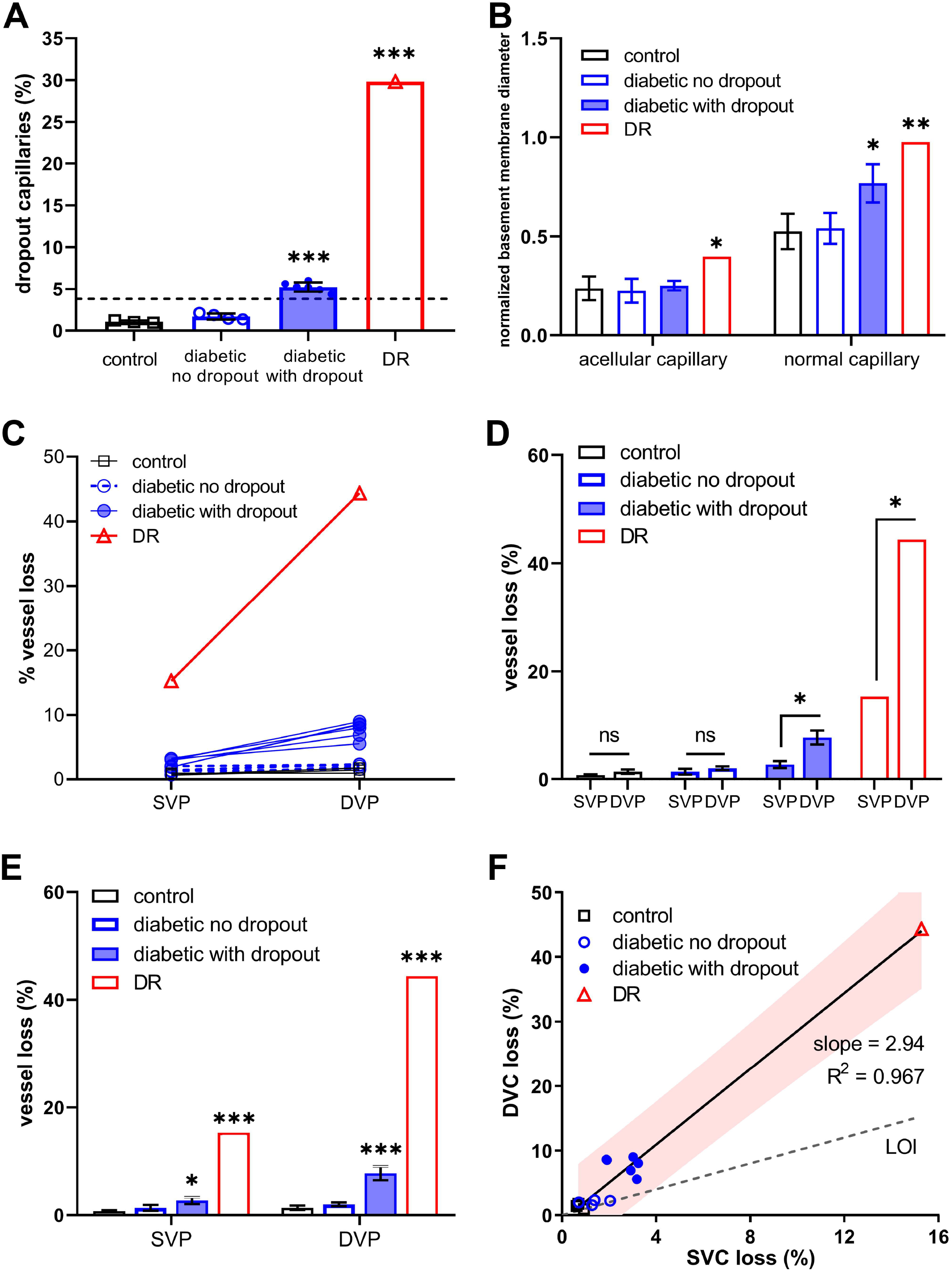
Vessel loss at different retinal vascular plexuses. **A**, Total capillary loss in control, subgroups of diabetic, and DR eyes. Diabetic retinae presented a mildly, but statistically significant, higher incidence than controls. DR retina presented an overwhelmingly higher dropout incidence than any other groups. The dashed line showed average capillary loss incidence in the diabetic group as a whole (3.82%). At least 13,500 vessels were assessed for each donor. **B**, Capillaries became narrower after losing endothelial cells. Slight increase in the size of normal capillaries was seen in the diabetic with dropout subgroup, but the size of acellular capillaries was consistent amongst non-DR groups. At least 15 capillaries were measured from each donor, no less than 52 capillaries were measured for each group. **C and D**, Intragroup differences in vessel loss of SVP and DVP. Plotting vessel loss from each donor revealed that SVP loss was never higher than that of DVP in all cases examined (**C**). On average, only diabetic with dropout subgroup and DR group had higher incidence of vessel loss (**D**). (At least 9,000 vessels were examined for each plexus for each donor.) **E**, Intergroup differences in vessel loss of SVP and DVP. Control and diabetic no dropout subgroup showed no inter-plexus difference, which was significant in diabetic with dropout subgroup and DR, with DVP showing considerably elevated proportion of vessel loss. **F**, A linear regression model plotting vessel loss in the DVP against SVP in all donors and revealed a strong linear relationship between (R^2^ = 0.972, p < 0.0001). Control and diabetic no dropout subgroup formed a clearly separate cluster from the diabetic with dropout subgroup. DR locates at a distinct region that is further away for the rest groups. N = 3 (control), 4 (diabetic no dropout), 6 (diabetic with dropout) and 1 (DR). Results were presented as mean ± s.d. Box and whisker show the mean ± s.d. Statistical significance was tested by one-way ANOVA with *post hoc* Dunnett’s analysis (**A, B, E**) or two-tailed Student’s t-test (**D**). ns denotes not significant, ^*^ P < 0.05, ^**^ P < 0.005, ^***^ P < 0.001. Abbreviations: LOI, line of identity; SVP, superficial vascular complex; DVP, deeper vascular complex; DR, diabetic retinopathy.

To investigate whether acellularity leads to morphological changes of the vascular basement membrane, we measured the inner diameter of acellular and normal capillary basement membranes. Data was normalized to the size of RPE nuclei of each donor to correct for potential tissue shrinkage during processing. Results showed that the capillary basement membrane profile shrinks in diameter (by about 50%) after endothelial cells loss in retinas from all donor groups (**Figure 3B**). Furthermore, a mild dilation of normal capillary profiles in the DDO subgroup was observed (1.46-fold, p = 0.0051), whilst the diameter of the acellular capillaries remained comparative to controls (p = 0.72). In addition, the DR eye presented an almost two-fold dilation in both acellular (1.67-fold, p = 0.457) and normal capillaries (1.85-fold, p = 0.0046) compared to control.

### Vessel loss is more severe in the deep capillary plexus

It is worth noting that in the DNDO subgroup, no obvious vascular lesions were detected on the whole mount. This may be because in the whole mount images the DVP is more difficult to see than SVP. We therefore assessed in our analysis on cross sections the DVP and SVP separately. In all donors examined, we found that vessel loss in the DVP was always more pronounced than in the SVP (**Figure 3C**). On average, differences in inter-plexus vessel loss was not significant in controls (1.38% *vs* 0.75%, p = 0.07) or in the DNDO subgroup (2.02% *vs* 1.36%, p = 0.08), but was significant in the DDO subgroup (7.77% *vs* 2.67%, p < 0.0001) and the case with DR (15.28% vs 44.38%, p < 0.001) (**Figure 3D**). Inter-group differences of vessel loss were not significant between controls and the DNDO subgroup (p = 0.13 in SVP, p = 0.07 in DVP). But there was a significant increase in the DDO subgroup (p < 0.001 in both plexuses), with a steeper increase observed in the DVP. Vessel loss in the DR case was always significantly higher in both plexuses when compared with any other group at any plexus (p < 0.001) (**Figure 3E**).

Since we found that vessel loss in the DVP is always higher than in the SVP, we asked whether there was a potential correlation between them. **Figure 3F** reveals that there is a linear positive association between the DVP and the SVP (R^2^ = 0.972, p < 0.0001). We could identify a cluster formed by controls and the DNDO subgroup in Fig. 3F (bottom left plot), indicating low level of capillary dropout in both plexuses, from which the DDO subgroup forms a clearly separate cluster. Further away at the top right corner is the DR case, with a considerably higher level of vessel loss in both plexuses. All data were within the 99% confidence interval area (**Figure 3F**, red region) from the best fit model (**Figure 3F**, solid line), with the formula y=2.94x-0.84. This means with every 1% increase of vessel loss at the SVP, loss in the DVP will increase by around 3%.

### Effect of vascular dropout on gliovascular unit

To test how the integrity of the gliovascular unit is affected by the capillary dropout during the earliest stages of DR, we studied sections from the DDO subgroup by immunohistochemistry using antibodies against glial markers, aquaporin 4 (AQP4), cellular retinaldehyde–binding protein (CRALBP), glutamine synthetase (GS) and glial fibrillary acidic protein (GFAP).

GFAP expression was usually confined to retinal astrocytes and was either not upregulated or predominantly affected by microvascular dropout in eyes with diabetes (**Figure 4A**). GS was highly expressed in Müller cell endfeet at ILM and OLM, and processes in the OPL which were not tightly associated with vessels (**Figure 4B**). CRALBP was found in the Müller cell soma, processes in the OPL and the RPE (**Figure 4C**). CRALBP expression around Müller cell endfeet was visible in both normal (**Figure 4C**, zoomed in images, arrows) and acellular capillaries (**Figure 4C**, zoomed in images, arrowheads), however no obvious difference in staining intensity could be noticed. AQP4 was expressed predominantly at the endfeet and closely associated with the gliovascular unit (**Figure 4D**). Although AQP4 staining could still be seen around nonperfused capillaries (**Figure 4D**, zoomed in images, arrowheads), the staining intensity was clearly lower than that around normal capillaries **Figure 4D**, zoomed in images, arrows).

**Figure 4.**
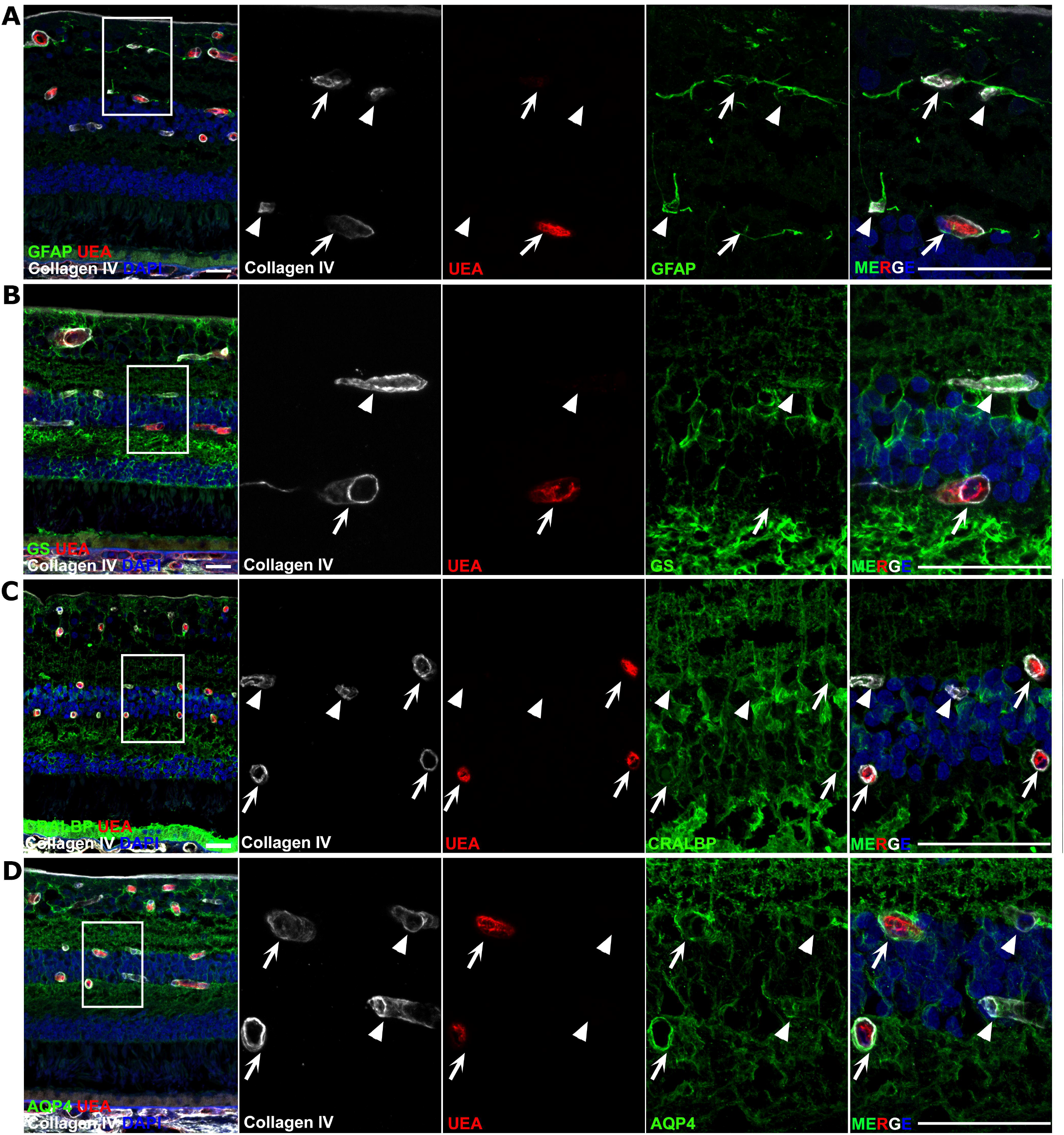
Gliovascular Units in diabetic retina. Immunohistochemistry using antibodies against widely recognised glial marker GFAP (**A**), GS (**B**), CRALBP (**C**) and AQP4 (**D**) from consecutive sections of donors from diabetic with dropout subgroup. Arrowheads point at acellular capillaries and arrows show normal capillaries. Glial interface formed by retinal astrocytes was intact regardless of vascular dropout (**A**). GS and CRALBP are highly expressed in Müller cell endfeet at the ILM and OLM, and less involved in the gliovascular interface (**B** and **C**). AQP4 expression is specially concentrated around glial cell endfeet around vessels, which remained present around acellular capillaries (**D**, arrows), but staining intensity was notably reduced. (**D**, arrowheads). Scale bar 50 μm in lower magnified images and 25 μm in zoom-in images.

### Deep capillary’s zone of influence

Next, we attempted to evaluate the local effect of capillary dropout (and presumed loss of oxygen supply) on the survival of neural cells, especially neurons in the vicinity of acellular DVP capillaries. To this end we first determined the zone of influence of capillaries in the DVP by measuring the nearest neighbour distance (NND, distance between neighbouring capillaries), which was constant across groups (**Figure 5A**). Furthermore, distribution analysis of NND revealed consistency between control and diabetic subgroups. Although the peak of DR NND did not shift, it was clearly lower than other groups (**Figure 5B**). This can be further described by the regularity index (RI), defined as the ratio of the mean NND over its standard deviation. Despite of consistent NND across all groups, the RI of the DR case showed statistically significant RI reduction (p < 0.05); whereas there was no difference amongst control and diabetic subgroups (**Figure 5C**). To assess whether NND changes with respect to retinal eccentricity, multiple sections with varying distance to the optic disc-fovea axis were analysed for each group. As shown in **Figure 5D**, NND plotted against the distance to the optic disc-fovea axis showed no changes in the superior-inferior axis (DR and diabetic no dropout subgroup not shown).

**Figure 5.**
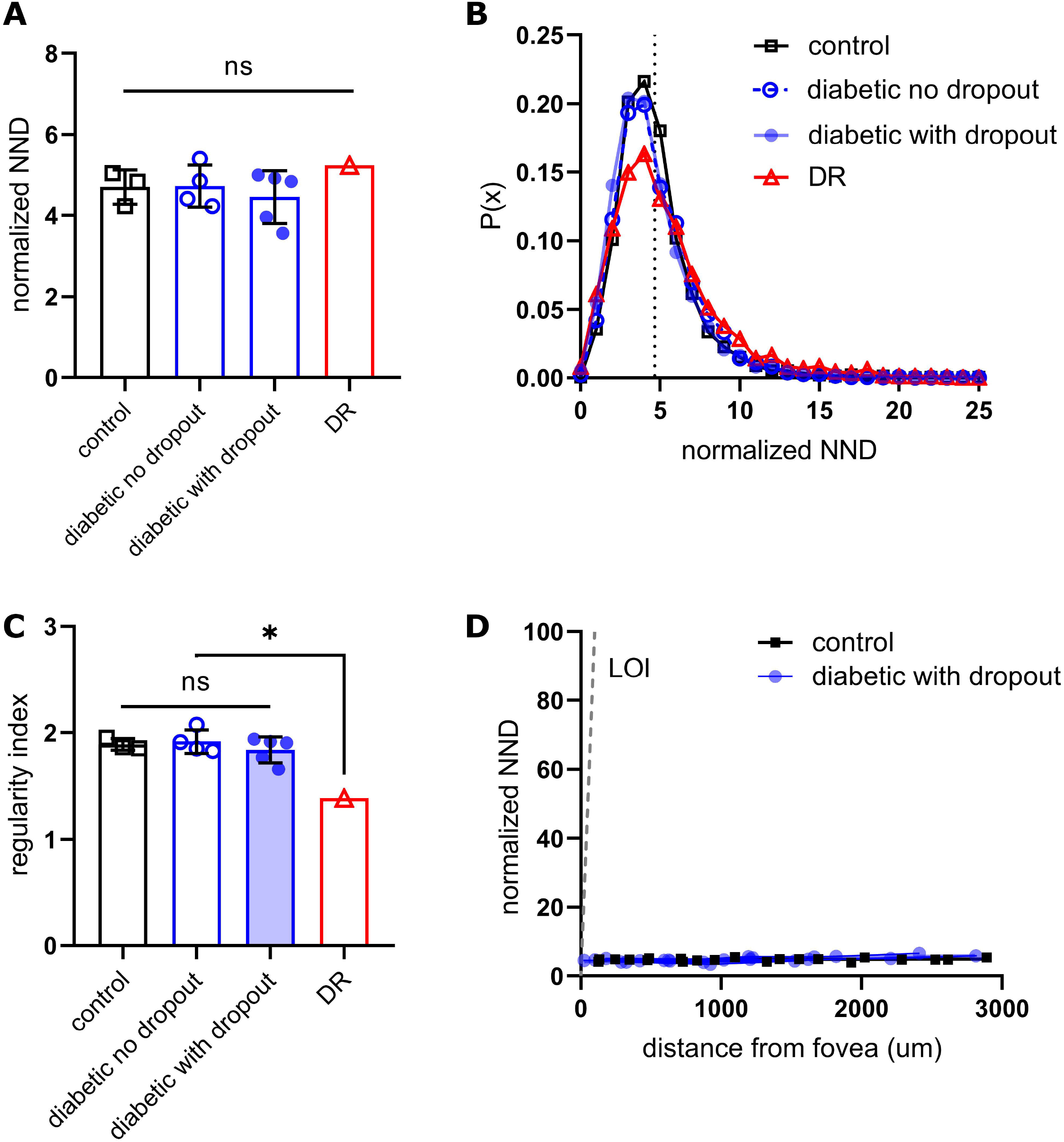
DVP capillary zone of influence. **A**, NND was calculated for each donor and then normalized to the height of the nuclei of RPE cells (over 10 RPE cells were measured per donor and the average value was used), so that results express fold change to the reference value. No difference was found between any two groups, meaning that the NND is consistent across groups. **B**, Frequency plot of the NND reveals an overlapping distribution pattern in control and two diabetic subgroups, while lower peak of the DR NND distribution was noticed. The dashed line shows the overall normalized mean NND of 4.65. **C**, Regularity index, calculated as mean/s.d. tends to decrease with increased severity of capillary loss. No difference was found amongst control and subgroups of diabetics, whereas RI of DR was significantly different from every one another. **D**, The NND was plotted against the distance from the optic disc-fovea axis along the superior-inferior axis. No linear relationship could be found in any group (diabetic no dropout subgroup and DR not shown). N = 3 (control), 4 (diabetic no dropout), 5 (diabetic with dropout), 1 (DR). Results are presented as mean ± s.d., n.s. denotes not significant and ^*^P < 0.05. Statistical significance was tested by one-way ANOVA with Tukey’s post hoc comparison (A, C). Abbreviations: LOI, line of identity; NND, nearest neighbour distance.

### Neural loss in the INL irrespective of DVP vascular dropout

Having determined the NND as a definition of DVP zone of influence, we quantified the number of nuclei within a circular area (with the NND x 0.5 as its radius) from each capillary in the DVP. An example is shown in **Figure 6A**. Results showed a comparable number of cells around normal and nonperfused capillaries in diabetic with dropout subgroup (12.27 ± 1.80 vs 12.39 ± 1.65, p = 0.9994) and DR (8.42 ± 2.33 vs 8.35 ± 1.33, p > 0.999). However, compared with controls, there was a general loss of 7% INL cells in the DDO subgroup (13.25 ± 2.27 vs 12.33 ± 1.71, p = 0.0457), which increased to 37% in DR (13.25 ± 2.27 vs 8.38 ± 1.89, p < 0.0001). Taken together, these results suggest that in patients with diabetes and no DR, there is a subtle but statistically significant loss of INL cells, which was much more pronounced in the DR case. Importantly, this effect was pan-retinal and independent of local capillary dropout (**Figure 6B**).

**Figure 6.**
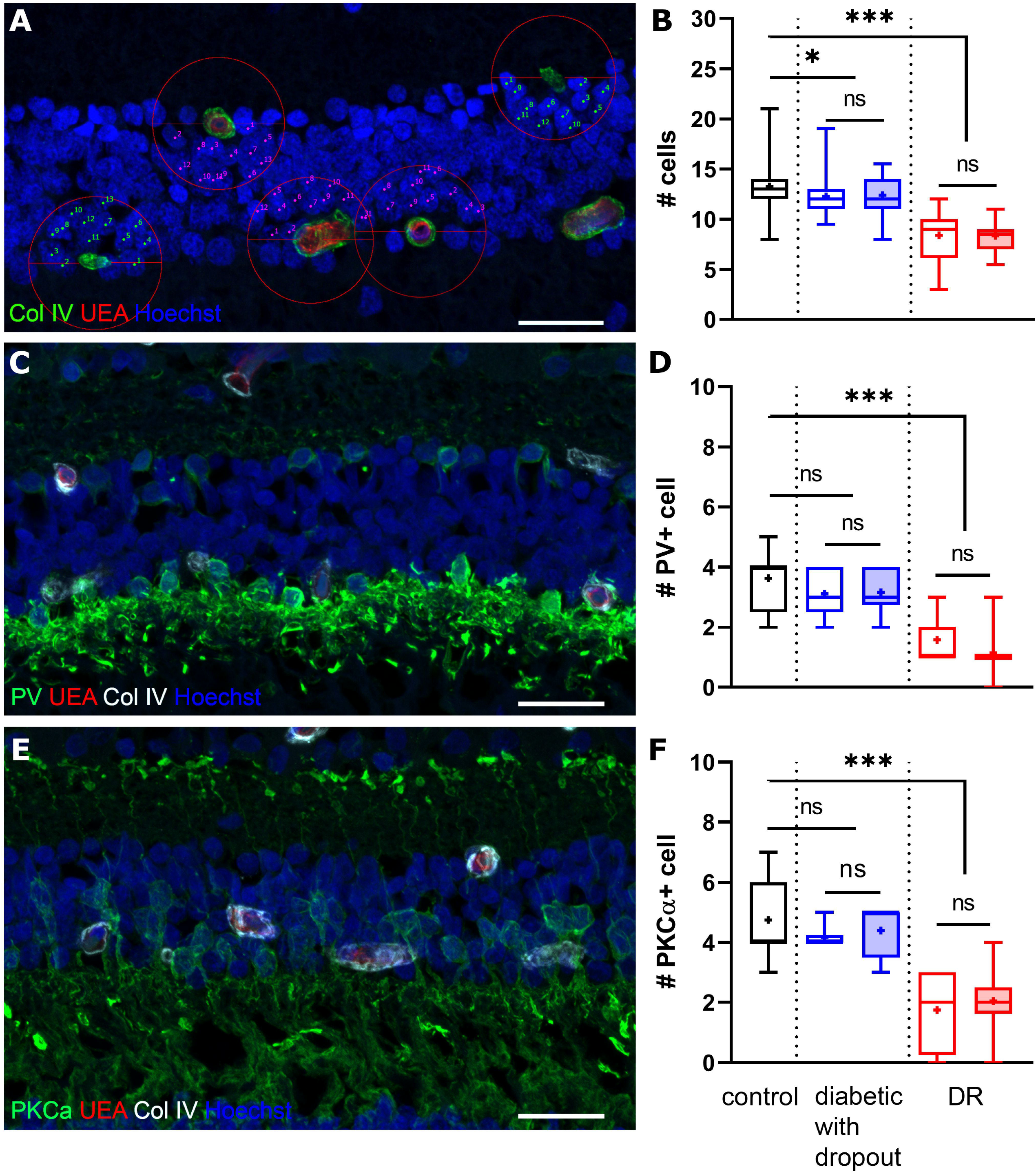
Identifying overall and subpopulation cell loss in the INL. A, C and E, confocal microscopy image showing the measurements of INL cell nuclei within the zone of influence of DVP capillaries (circle, A). Examples of normal capillaries (magenta counting marks) and nonperfused capillaries (green counting marks) are shown. Horizontal (C) and bipolar celIs (E) were visualized using antibodies against PV and PKCα, respectively. B, D and F, quantification of total cells (B) and interneurons (D and F) within deeper capillary zones of influence. Normal capillaries are shown by empty boxes, nonperfused capillaries are shown by filled boxes. N = 3 (control), 4 (diabetic no dropout), 5 (diabetic with dropout), 1 (DR). Box represents 25th, median and 75th quartile, “+” shows the mean value. Whiskers represent max and min. n.s. denotes not significant and ^*^ P < 0.05, ^***^ P < 0.0001. Statistical significance was tested by unpaired two-tailed Student’s test (intragroup differences) or one-way ANOVA with Dunnett’s post hoc comparison with control (intergroup differences). Scale bar 25 μm

Cells that contribute to the ERG b-wave may be affected in patients with diabetes and rodent models with diabetes. We therefore used antibodies raised against PV and PKCα to label horizontal (**Figure 6C**) and bipolar cells (**Figure 6E**), respectively. Quantifying these interneurons within deeper capillary zones of influence showed results in line with our previous finding that cell loss is independent of local capillary dropout (**Figure 6 D and F**). Of note, in the DDO subgroup, a subtle loss of around 10% PV+ horizontal cells (3.63 ± 1.07 vs 3.13 ± 0.74, p = 0.31) and PKCα+ bipolar cells (4.75 ± 1.39 vs 4.27 ± 0.65, p = 0.51) compared controls was identified. Although this difference was not statistically significant, it represents an interesting trend, because this cell loss was much more profound in the DR case, where around 60% of both interneuron types were lost compared to that in control (PV+ cells: 3.63 ± 1.07 vs 1.36 ± 0.84; PKCα+ cells: 4.75 ± 1.39 vs 1.97 ± 1.09; p < 0.0001 for both) (**Figure 6 D and F**).

## DISCUSSION

In this study, we have established capillary loss in the deeper plexus of the retinal vasculature as a novel and sensitive histological biomarker for the very early stages of DR in human postmortem tissue. This early sign of DR is likely to occur before a clinical diagnosis (typically based on visible changes in the SVP) and was remarkably common (6 out of 10) in our randomly selected group of tissue donors with diabetes.

Perfusion of the DVP and SVP has been extensively studied in recent years in living diabetics using OCTA (reviewed in supplemental material 1). Most of these studies showed that perfusion deficiencies were more pronounces in the DVP and several of them found such changes in people with diabetes and no DR signs, which aligns with our histological findings. Other histopathological studies have also described a higher incidence of more advanced vascular pathology, such as microaneurysms within the INL [28, 29].

One possible explanation for the increased occurrence of vascular damage in the DVP is that these vessels are more distal than vessels in the SVP. A small perfusion dysfunction in the SVP may have a more profound impact on the downstream vessels in the DVP (all of which receive their supply from the SVP). An alternative explanation might be that the only glia component contributing to gliovascular unit in the DVP comes from Müller cells, whereas in the SVP retinal astrocytes are also present. In our study we have shown altered distribution of AQP4 in Müller cell endfeet, possibly leading to water homeostasis defects in DR. This may be related to a compromised blood-retina barrier associated with DR [30], contributing to vascular leakage and oedema. However, in our tissue cohort with diabetes, immunohistochemistry with antibodies against human IgG did not reveal any signs of serum leakage into the retina (data not shown).

Another widely described feature of DR is increased vessel diameter. This matches our finding of wider, perfused capillaries in the early DR group and the DR case. OCTA measurements in patients have also shown increased vessel calibres in DR [31, 32]. The calibre increase we observed may be explained by a thickening of vessel basement membrane, as in diabetic animal models [33, 34]. Alternatively, loss of pericytes described in human diabetic/DR eyes may lead to vessel dilation [3, 29].

Our data also identifies neural cell loss in the INL as an early neurodegenerative change, which could be a consequence of reduced perfusion of the DVP. Oxygen tension in the INL is low [35] and the DVP may be an evolutionary adaptation to provide the INL with oxygen. Perfusion defects in the DVP may lead to excessive hypoxia and eventually cell loss in INL. Remarkably, even at the very early stage of vascular pathology found in our cases, we detected retinal neural cell loss, suggesting that vascular and neural defects are closely intertwined at the earliest stages of DR. Interestingly, baseline DVP nonperfusion (assessed by OCTA) in NPDR patients can predict DR progression with high accuracy [36, 37]. Moreover, it was shown that the parafoveal vessel density in the DCP is the parameter most robustly associated with the clinical stage of nonproliferative DR [38]. This not only highlights the potential use case of microvascular changes in the DVP as a biomarker to predict the DR progression, but it also indicates a potential functional interaction.

However, the lack of spatial correlation between localised capillary and neural loss observed in our study seems to exclude the simple mechanism of localised hypoxia causing neural cell apoptosis in the vicinity of acellular capillaries. Nevertheless, it cannot be excluded that vascular dysfunction (currently not detectable in postmortem tissue), such as a lack of perfusion autoregulation, may cause the diffuse cell loss in the INL we observed. Alternatively, diabetes may have a direct impact on neural cells in the retina independently of - or possibly even before - vascular damage [39–42]. As non-invasive in vivo clinical imaging improves [43], longitudinal studies in people with diabetes may provide further insights about the temporal relationship between vascular and neural dysfunction.

In summary, our findings contribute to the DR literature showing strong evidence for early retinotopic vascular and neurodegenerative changes which are of clinical importance and are meaningful for early DR detection. DVP changes and INL atrophy might be incorporated into newer DR classification systems to reap the benefits of early diagnosis and strengthening measures to improve metabolic control in patients with early DR changes.

## ACKNOWLEDGEMENTS

The authors thank Meaghan O’Neill for technical assistance and Almas Dawood for help with initial processing of the diabetic retinopathy eye.

## Funding

Diabetes UK, Santen Pharmaceutical PhD studentship, NIHR Moorfields Biomedical Research Centre

MF, CE and AT conceived and designed the analysis; QY, JM and MY collected the data; QY, AOB, JM and MY contributed data or analysis tools; QY performed the analysis; QY and MF wrote the manuscript.

## FIGURE LEGENDS

**Supplementary material 1. Review of clinical OCTA studies investigating vascular nonperfusion in diabetes and DR**.

Anatomical abbreviations: SCP, superficial capillary plexus; DCP, deep capillary plexus; CC, choriocapillaris; FAZ, foveal avascular zone; DR, diabetic retinopathy; NDR, diabetic with no DR; T1DM, type 1 diabetes mellitus; T2DM, type 2 diabetes mellitus; T1/2DM, type 1 or 2 diabetes mellitus; ILM, inner limiting membrane; IPL/INL, inner plexiform layer / inner nuclear layer interface; INL/OPL, inner nuclear layer / outer plexiform layer interface; OPL/ONL, outer plexiform layer / outer nuclear layer interface. Parameter abbreviations: VD, vessel density; PAN, percent area of nonperfusion; AFI, adjusted flow index; CPD, capillary perfusion density; FD, flow density; VAD, vessel area density; VLD, vessel length density; VDI, vessel diameter index.

**Supplementary material 2. 3D image of a ghost vessel from diabetic tissue**.

Immunostaining using UEA (red) and antibody against collagen IV (green) on a piece of retina from a diabetic eye in the with-dropout-group. The video shows a 3D view of a ghost vessel in the superficial plexus. The ghost vessel appears pure green (on the righthand side of the imaged sample). The green/yellow cells inside vessels are auto fluorescent erythrocytes.

