## Supplementary Material 1 for "Disconnection Between Microvascular Damage and Neurodegeneration in Early Diabetic Retinopathy"

**Supplementary Material 1. Clinical OCTA studies investigating vascular nonperfusion in DR.**

| Participants (eyes) | Scan size  (mm x mm) | OCTA device | Segmentation criteria | Key findings | Reference |
| --- | --- | --- | --- | --- | --- |
| Nonperfusion in DCP > SCP (12 papers) | | | | | |
| 84 T2DM any DR,  34 controls | 3 × 3 | RTVue-XR Avanti (Optovue, USA) | SVP: ILM TO 110μm above RPE | VAD and VLD are notably lower only in DCP between mild NPDR and moderate-severe NPDR/PDR. | [1] |
|  |  |  | DVP: 110μm above RPE to RPE |  |  |
| 25 T1DM NDR,  25 controls | 3 × 3 | Cirrus 5000 (Carl Zeiss Meditec, Germany) | Not specified | Decreased DVP density in NDR eyes, no difference in SVP or CC density. | [2] |
| 47 T1/2DM DR,  29 controls | 3 × 3 and  6 × 6 | RTVue-XR Avanti (Optovue, USA) | SVP: 3 to 15μm from ILM | FD is significantly reduced in DVP in NDR compared to SVP. | [3] |
|  |  |  | DVP: 15 to 70μm from ILM |  |  |
| 28 T1DM NDR or mild DR,  23 controls | 3 × 3 | RTVue-XR Avanti (Optovue, USA) | SVP: ILM to IPL | Reduction in parafoveal DCP density in T1DM patients with no or mild signs of DR. | [4] |
|  |  |  | DVP: IPL to OPL |  |  |
| 33 T2DM NDR,  29 controls | 3 × 3 | RTVue-XR Avanti (Optovue, USA) | SVP: 3 to 15μm from ILM | Statistically significant reduction in SCP and DVP vascular density in NDR compared to control, especially in DVP. | [5] |
|  |  |  | DVP: 15 to 70μm from ILM |  |  |
| 102 T1/2DM DR,  62 controls | 6 × 6 | RTVue-XR Avanti (Optovue, USA) | SVP: ILM to IPL/INL | Reduction in perfusion indices was significantly pronounced in DVP than SVP in the perifovea. | [6] |
|  |  |  | DVP: IPL/INL to OPL/ONL |  |  |
| 20 T1DM NDR,  23 controls | 3 × 3 | AngioVue OCTA (Optovue, USA) | SVP: ILM to IPL/INL | Reduced vessel density in the DVP in NDR. | [7] |
|  |  |  | DVP: IPL/INL to OPL/ONL |  |  |
| 71 T2DM NDR,  67 controls | 6 × 6 | RTVue-XR Avanti (Optovue, USA) | SVP: 3 to 15μm from ILM | Reduced vessel density in SVP and DVP in T2DM NDR patients, in which DCP is affected more. No difference in FAZ area. | [8] |
|  |  |  | DVP: 15 to 70μm from ILM |  |  |
| 22 T1D DR,  12 controls | 3 × 3 and  6 × 6 | RTVue-XR Avanti (Optovue, USA) | SVP: 3μm below ILM to IPL | Vascular density is reduced in diabetic eyes with lower visual acuity than in those with normal visual acuity in all vascular plexuses. Visual acuity is associated with degree of capillary loss in the DVP. | [9] |
|  |  |  | IVP: IPL to 9μm above OPL |  |  |
|  |  |  | DVP: 19μm below IN;/OPL to 9μm below OPL/ONL |  |  |
| 102 varying DR,  30 NDR,  42 controls | 3 × 3 | DRI OCT Triton plus (Topcon, Japan) | SVP: 2.6μm below ILM to 15.6μm below IPL/INL | FAZ area and perimeter correlate positively with DR severity. Decreasing trend of FAZ CI at DVP. Retinal microvasculature changes in DVP precedes that in SVP. | [10] |
|  |  |  | DVP: 15.6μm below IPL/INL junction to 70.2μm below IPL/INL |  |  |
| 60 T1/2DM NDR,  30 controls | 3 × 3 | DRI OCT Triton plus (Topcon, Italy) | SVP: ILM to 15.6μm above IPL/INL | Almost all NDR patients presented parafoveal capillary loss, with higher incidence in the DVC. | [11] |
|  |  |  | DVP: 15.6μm above to 70.2μm below IPL/INL |  |  |
| 396 eyes with various severity of DR | 3 × 3 | RTVue-XR Avanti (Optovue, USA). | SVP: 3μm below ILM to IPL/INL | Vascular density of all layers reduces with increased DR severity. In eyes with no to early DR, vascular changes in the DVP are most prominent, however the opposite in eyes with advanced DR. | [12] |
|  |  |  | IVP: IPL/INL to 20μm below IPL/INL |  |  |
|  |  |  | DVP: 20μm below IPL/INL to 15 μm below OPL/ONL |  |  |
| Nonperfusion in SCP > DCP (4 papers) | | | | | |
| 18 T1/2DM DR,  22 controls | 3 × 3 | DRI OCT Triton (Topcon Corp., Japan) | SVP: ILM to IPL/INL | Significantly reduced SVP vascular density in mild and moderate NPDR compared to controls; differences in DVP were not statistically significant. | [13] |
|  |  |  | DVP: IPL/INL to INL/OPL |  |  |
| 92 any DR,  44 NDR,  44 controls | 3 × 3 | RTVue-XR Avanti (Optovue, USA) | SVP: 3μm below ILM to 25um above IPL | In all three plexuses, with worsening DR, vascular density decreases while PAN increases. | [14] |
|  |  |  | IVP: IPL/INL to 30μm below IPL |  |  |
|  |  |  | DVP: 15μm slab below INL |  |  |
| 86 any DR,  44 controls | 3 × 3 | RTVue-XR Avanti (Optovue, USA) | SVP: 3um below ILM to 15um below the IPL | PAN and AFI is positively and negatively related to DR severity, respectively. DVP vascular density correlates strongly with DR severity. | [15] |
|  |  |  | DVP: 15 to 70um below IPL |  |  |
| 84 any DR,  14 controls | 3 × 3 | Cirrus SD-OCT (Carl Zeiss Meditec, USA) | SVP: ILM to 110um above RPE | Statistically significant reduction in SD, vascular density and FD, and increase in VDI, between healthy and moderate-severe NPDR and PDR in both plexuses. | [16] |
|  |  |  | DVP: 110um above RPE to RPE |  |  |
| Did not compare between plexuses (4 papers) | | | | | |
| 56 varying DR,  21 controls | 3 × 3 and  6 × 6 | RTVue-XR Avanti (Optovue, USA | SVP: ILM to IPL/INL | CPD values significantly lower in nearly all layers of all DR groups compared with control. | [17] |
|  |  |  | DVP: IPL/INL to OPL/ONL |  |  |
| 81 T2DM DR,  19 T2DM NDR | 3 × 3 | Swept-source OCTA (Topcon Corp., Japan) | SVP: 3 to 15μm from ILM | FD in both SCP and DCP is positively related to DR severity. No comparisons made between plexuses. | [18] |
|  |  |  | DVP: 15 to 70μm from ILM |  |  |
| 17 T1DM with severe NPDR or PDR,  17 controls | 3 × 3 | RTVue-XR Avanti (Optovue, USA) | SVP: ILM to 9μm above IPL/INL | Vascular density decreases significantly with DR severity in all three vascular plexuses. Inner retinal thickness correlated with vascular density in the SVP, but outer retinal thickness does not correlate with DVC vascular density. | [19] |
|  |  |  | IVP: 9μm above IPL/INL to 6μm below INL/OPL |  |  |
|  |  |  | DVP: 6μm below INL/OPL to 9μm below OPL/ONL. |  |  |
| 24 T1DM NDR,  24 controls | Not specified | RTVue-XR Avanti (Optovue, USA) | SVP: 3μm from ILM to IPL/INL | Attenuation of both SVP and DVP in the T1DM group compared with the control group in para and perifoveal regions. | [20] |
|  |  |  | IVP: IPL/INL to 20μm below IPL/INL |  |  |
|  |  |  | DVP: 20μm below IPL/INL to 15μm below OPL/ONL. |  |  |

Abbreviations: SVP, superficial vascular plexus; DVP, deep vascular plexus; CC, choriocapillaris; FAZ, foveal avascular zone; DR, diabetic retinopathy; NDR, diabetic with no DR; T1DM, type 1 diabetes mellitus; T2DM, type 2 diabetes mellitus; T1/2DM, type 1 or 2 diabetes mellitus; ILM, inner limiting membrane; IPL/INL, inner plexiform layer / inner nuclear layer interface; INL/OPL, inner nuclear layer / outer plexiform layer interface; OPL/ONL, outer plexiform layer / outer nuclear layer interface.

Parameter abbreviations; PAN, percent area of nonperfusion; AFI, adjusted flow index; CPD, capillary perfusion density; FD, flow density; VAD, vessel area density; VLD, vessel length density; VDI, vessel diameter index.
